# Multiscale simulations reveal key features of the proton pumping mechanism in cytochrome *c* oxidase

**DOI:** 10.1101/040717

**Authors:** Ruibin Liang, Jessica M. J. Swanson, Yuxing Peng, Mårten Wikström, Gregory A. Voth

**Affiliations:** Department of Chemistry, Institute for Biophysical Dynamics, James Franck Institute, and Computation Institute, University of Chicago, 5735 S. Ellis Ave., Chicago, Illinois 60637, USA; Research Computing Center, University of Chicago, 6030 S. Ellis Ave, Suite 126, Chicago Illinois 60637, USA; Helsinki Bioenergetics Group, Programme for Structural Biology and Biophysics, Institute of Biotechnology, University of Helsinki, FI-00014, Helsinki, Finland

## Abstract

Cytochrome *c* oxidase (C*c*O) reduces oxygen to water and uses the released free energy to pump protons across the membrane, contributing to the transmembrane proton electrochemical gradient that drives ATP synthesis. We have used multiscale reactive molecular dynamics simulations to explicitly characterize (with free energy profiles and calculated rates) the internal proton transport events that enable pumping and chemistry during the A→P_R_→F transition in the *aa*_3_-type C*c*O. Our results show that proton transport from amino acid residue E286 to both the pump loading site (PLS) and to the binuclear center (BNC) are thermodynamically driven by electron transfer from heme *a* to the BNC, but that the former (i.e., pumping) is kinetically favored while the latter (i.e., transfer of the chemical proton) is rate-limiting. The calculated rates are in quantitative agreement with experimental measurements. The back flow of the pumped proton from the PLS to E286 and from E286 to the inner side of membrane are prevented by the fast reprotonation of E286 through the D-channel and large free energy barriers for the back flow reactions. Proton transport from E286 to the PLS through the hydrophobic cavity (HC) and from D132 to E286 through the D-channel are found to be strongly coupled to dynamical hydration changes in the corresponding pathways. This work presents a comprehensive description of the key steps in the proton pumping mechanism in C*c*O.

**Significance:** The long studied proton pumping mechanism in cytochrome *c* oxidase (C*c*O) continues to be a source of debate. This work provides a comprehensive computational characterization of the internal proton transport dynamics, while explicitly including the role of Grotthuss proton shuttling, that lead to both pumping and catalysis. Focusing on the A to F transition, our results show that the transfer of both the pumped and chemical protons are thermodynamically driven by electron transfer, and explain how proton back leakage is avoided by kinetic gating. This work also explicitly characterizes the coupling of proton transport with hydration changes in the hydrophobic cavity and D-channel, thus advancing our understanding of proton transport in biomolecules in general.

## Introduction

Cytochrome *c* oxidase (C*c*O, Fig. 1) is the terminal enzyme in the respiratory electron transfer chain in the inner membrane of mitochondria and plasma membrane of bacteria. It catalyzes the reduction of O_2_ to H_2_O and couples the free energy of this exergonic reaction to the pumping of protons across the membrane, creating a transmembrane proton electrochemical gradient that drives, for example, ATP synthesis. During each reaction cycle eight protons are taken up from the negatively-charged inside (N-side) of the membrane and either react with oxygen (referred to as ‘chemical’ protons below) or are pumped to the positively-charged outside (P-side) of the membrane (referred to as ‘pumped’ protons below). In *aa*_3_-type C*c*O, as found in mitochondria, the D-channel is responsible for uptake of all four pumped protons and at least one out of four chemical protons. Protons on the N-side are taken into D-channel via the amino acid residue D132 at the channel entrance, and then transferred to residue E286 in the middle of the membrane. By Grotthuss shuttling through the water molecules in the hydrophobic cavity (HC) above E286, each proton is either transferred to react with oxygen in the binuclear center (BNC), consisting of heme *a*_3_ and the Cu_B_ complex, or transferred to the pump loading site (PLS) and then further released to the P-side of the membrane (c.f. Fig. 1). Despite decades of study, the C*c*O proton pumping mechanism, which entails the transport of two protons from the N-side of the membrane (one to be pumped and one for catalysis) coupled to a single electron transferred to the BNC, is still incompletely understood at the atomistic level. It has been unclear, for example, how electron transfer (ET) is coupled with the multiple proton transport (PT) events, in what order the charge transport processes happen, and how C*c*O prevents back flow of pumped protons during the transfer of chemical protons.

**Fig. 1.**
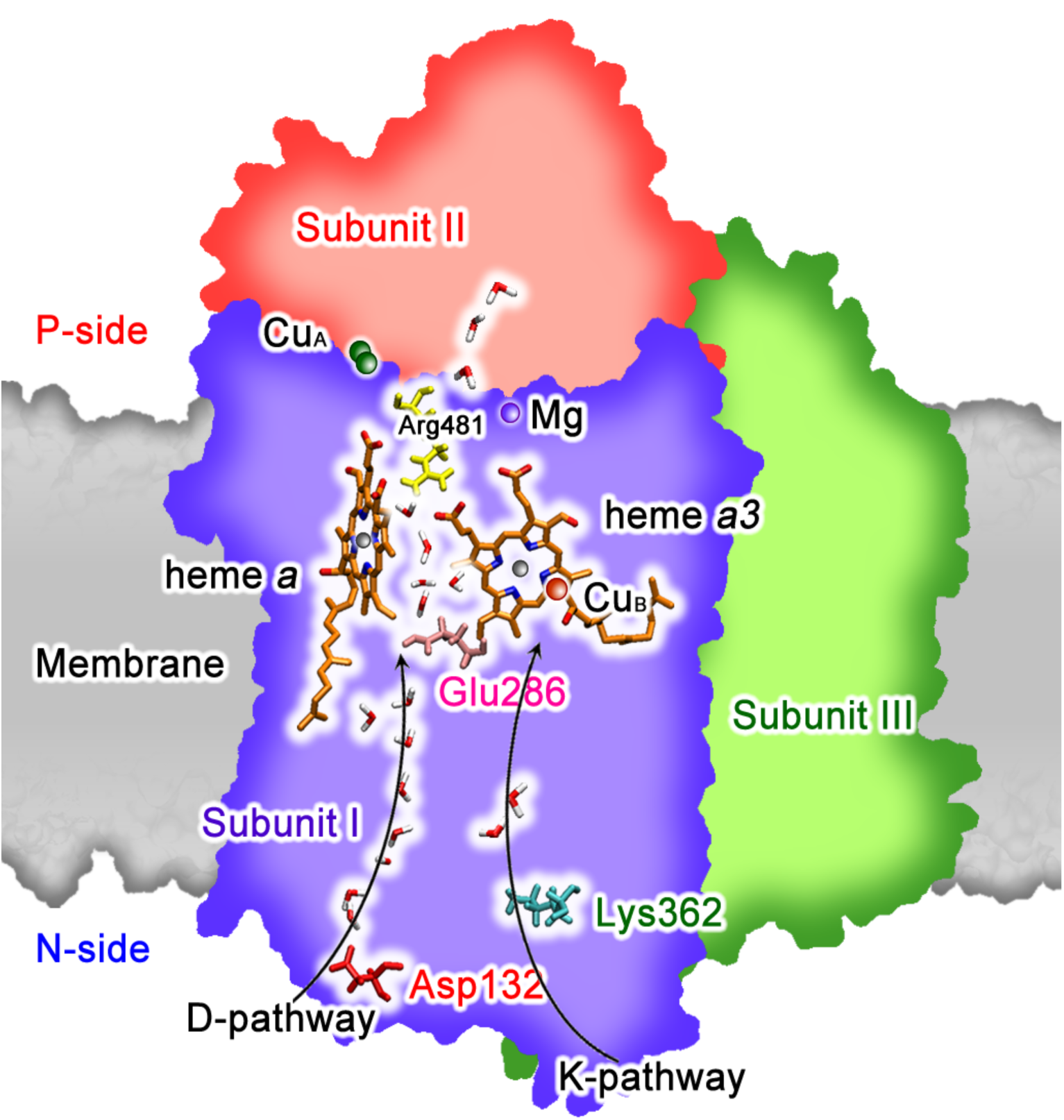
Illustration of the simulation setup for the full C*c*O from *Rhodobactor sphaeroides* in a membrane and surrounded by water. The D-and K-channels, as well as metal centers, key residues, and internal water molecules are depicted.

To address these questions, experimental results must be complemented with molecular-level insight from computer simulation. However, in large and complicated biomolecular systems it is challenging to simulate PT, which requires an explicit treatment of the positive charge defect associated with a hydrated excess proton, including its delocalization and Grotthuss shuttling (1). To overcome this challenge, a multiscale reactive molecular dynamics (MS-RMD) method has been extensively developed and applied in our group to study PT in aqueous and biological contexts [see, e.g., Refs. (2-7)]. Here we have carried out extensive MS-RMD free energy simulations to study the proton pumping mechanism in C*c*O. In our MS-RMD approach, quantum mechanical forces from targeted quantum mechanics/molecular mechanics (QM/MM) calculations are bridged, in a multiscale fashion via a variational mathematical framework, into the reactive MD algorithm (MS-RMD) for the dynamics of system nuclei, thus including chemical bond breaking and making. In this way, we explicitly simulate PT between proton binding sites, including Grotthuss shuttling of the excess proton(s) through residues and intervening water molecules.

We focus in this work on the A→P_R_→F transition in C*c*O, which occurs during the oxidation of the fully reduced enzyme, because there is extensive experimental information to which we can compare our computational results. Based on our simulations, we provide a quantitative, comprehensive and molecular-level description of proton uptake, pumping, and chemical proton transfer during the A→P_R_→F transition. Our results show that both PT events (E286 to the PLS and E286 to the BNC) are driven by ET from heme *a* to the BNC. The transfer of the pumped proton is kinetically favored while that of the chemical proton is rate limiting. Our calculated rate for the chemical proton transfer is in quantitative agreement with experimental measurements. (8, 9) These results also explain how C*c*O prevents the decoupling of pumping from the chemical reaction with kinetic gating. The fast pumping process precedes transfer of the chemical proton to the BNC, and fast D-channel PT to E286 after pumping prevents proton back flow from the PLS. Given the computational accuracy and efficiency of the MS-RMD methodology (7), a critical component of our results is the explicit characterization of the coupling between PT and hydration changes in both the HC and D-channel, revealing a remarkable and dynamic coupling between the migration of the excess proton and hydration. Finally, we present results that argue against the possibility of E286 being biprotonated during the pumping process.

## Results and Discussion

In the current work, three intermediate redox states during the A→P_R_→F transition were simulated: P_M_’, P_R_ and F (*SI Appendix*, Table S1). The first intermediate state is P_M_’, where both Cu_A_ and heme *a* are reduced, the BNC is oxidized, and Cu_B_ has a hydroxide ligand. Electron transfer from heme *a* to the tyrosine radical in the BNC converts P_M_’ into the P_R_ state. Following this, PT from E286 to the PLS and a second PT to the Cu_B_ bound hydroxide, forming a water molecule, converts P_R_ into F. Partial electron transfer from Cu_A_ to heme *a* and proton release to the P-side of the membrane, which we did not simulate in this work, completes the A→P_R_→F transition.

### Transport of the Pumped Proton and Hydration of the HC

Previous results have suggested that internal PT from E286 to the PLS and BNC are coupled to the redox states of heme *a* and the BNC (9-20). However, controversy remains regarding how they are coupled, in what order the PT and ET events occur, and what features enable pumping while preventing the back flow of protons from the P-side to the N-side (8, 9, 13-15, 19, 21). Particularly controversial has been the role of water in the HC during the proton pumping process with some authors arguing for a low hydration state (3-5 waters) (22-26), while others have suggested a high hydration state (> 6 waters) (27). Recent computational work proposed a stepwise pumping mechanism in which an excess proton is first transported from E286 to PRD*a*_3_ (the putative PLS) through a poorly hydrated HC, followed by an increase in the HC hydration (27), which was used to describe the transfer of the chemical proton (20). This issue is challenging to resolve because, as recently reported in ref (28), the migration of a hydrated excess proton can be strongly coupled with a dynamically changing solvation environment, to the extent that protons can even create their own “water wires” in otherwise dry hydrophobic spaces. In other words, as shown herein for C*c*O the two processes (PT and dynamic hydration) happen cooperatively, with waters entering and leaving the HC during the charge migration processes, and with the water hydration being intrinsically coupled to the proton charge defect translocation. Capturing this type of cooperativity often requires computationally demanding enhanced sampling of multiple degrees of freedom. The MS-RMD approach has allowed us to overcome this challenge for C*c*O.

To address the above mentioned controversies, we have simulated PT to the PLS and the BNC focusing on the coupled hydration changes. Starting with PT from E286 to the PLS we calculated *two-dimensional* free energy profiles, or 2D potentials of mean force (2D PMFs), in the P_M_’ (before ET) and P_R_ (after ET) states during the A**→** F transition (*SI Appendix*, Table S1). The collective variables used to define these 2D PMFs are (1) the progress of the excess proton center of excess charge (CEC) through the HC (horizontal axis) and (2) the degree of hydration of the HC (vertical axis). (see *SI Appendix* for definitions and more discussion). The 2D PMFs (Figs. 2 A and B) and minimum free energy pathways (black lines) verify that as the proton moves from E286 to the D-propionate on heme *a*_3_ (PRD*a*_3_) during the corresponding activated rate processes described by these pathways, the HC becomes more hydrated (increasing from six to ten waters in our chosen hydration “box”, which also includes approximately two waters outside of the traditionally defined inter-heme region). The curvy and non-horizontal nature of the minimum free energy pathways indicates that the two processes are indeed cooperative and coupled. When E286 is protonated, the HC favors a low hydration state (~ four waters in the inter-heme region) (*SI Appendix*, Fig. S5 A). As the excess proton moves to the water above E286 (*SI Appendix*, Fig. S5 B) the PRD*a*_3_ side chain rotates down to interact with the positive charge (CEC). This weakens the interactions between PRD*a*_3_ and nearby W172 and R481, allowing more water molecules to enter the HC. The hydration level in HC reaches its maximum just before (P_R_) or as (P_M_’) PRD*a*_3_ is protonated (*SI Appendix*, Fig. S5 C). The PRD*a*_3_ then rotates from downward to upward orientation through the transition state (*SI Appendix*, Fig. S5 D). Subsequently, the proton moves to the A-propionate on heme *a*_3_ (PRA*a*_3_, the final PLS identified in this work as discussed below), and the high hydration state remains stable as long as E286 remains deprotonated (*SI Appendix*, Fig. S5 E). (This behavior was further confirmed by 30 ns of classical simulations of the P_M_’ and P_R_ states with both E286 and PRD*a*_3_ deprotonated and PRA*a*_3_ protonated.)

**Fig. 2.**
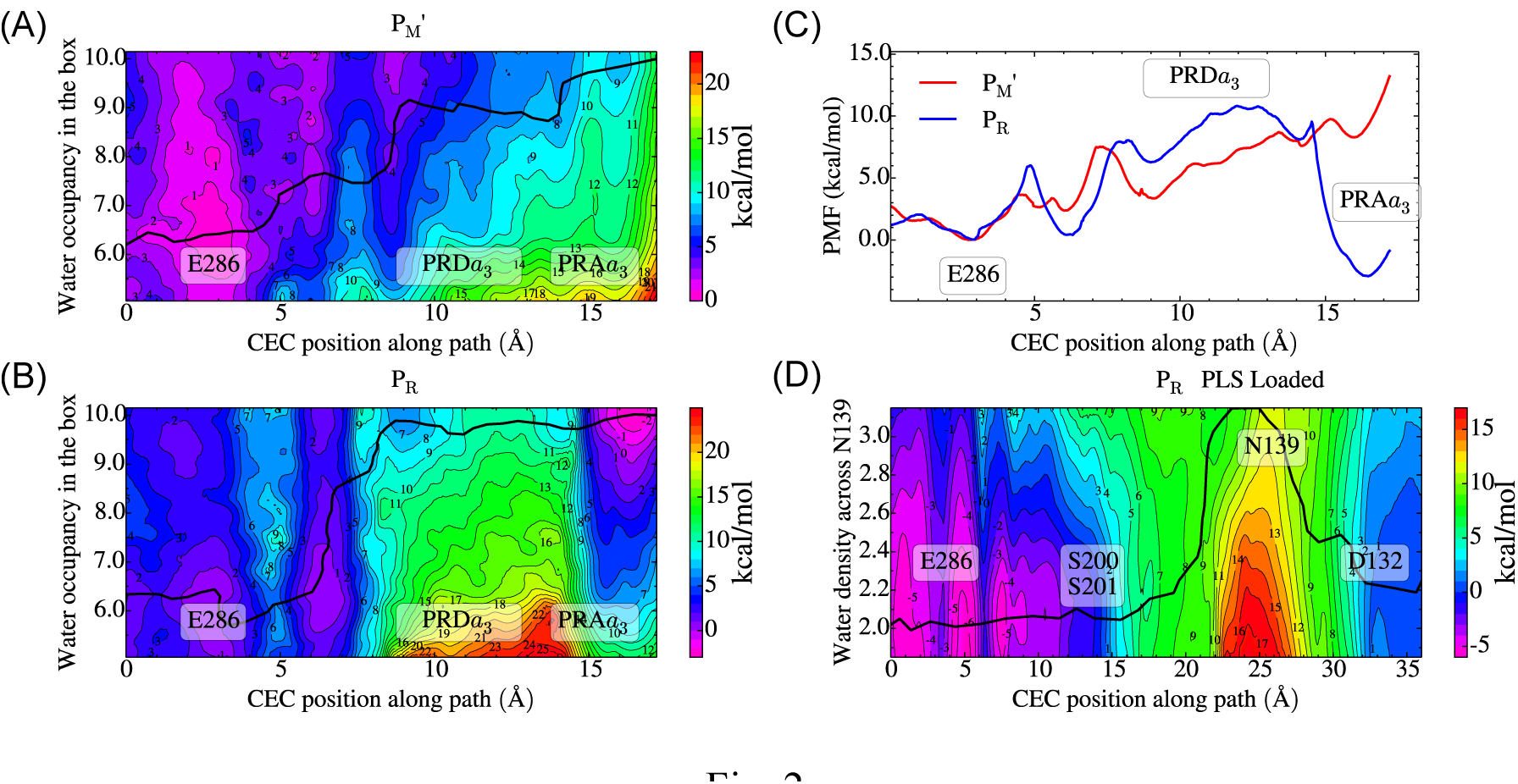
(A) and (B): Two-dimensional free energy profiles (2D-PMFs) for PT from singly protonated E286 to the deprotonated PLS in the P_M_’ and P_R_ states, respectively, as a function of the excess proton center of excess charge (CEC) coordinate through the hydrophobic cavity (HC) as the horizontal axis and the water hydration in the HC as the vertical axis. The minimum free energy pathways (black lines) are diagonal in nature, indicating the two processes are coupled. (C) 1D free energy profiles (PMFs) for PT in the HC along the minimum free energy pathway for the P_M_’ (red) and P_R_ (blue) states. (D) 2D-PMF for PT from singly protonated D132 to the deprotonated E286 through the D-channel in the P_R_ state, with proton preloaded at PLS. The 2D-PMF is a function of the CEC coordinate through the D-channel as the horizontal axis and the water hydration in the asparagine gate region as the vertical axis. The minimum free energy pathway is depicted by a black line. The strongly coupled behavior of the PT CEC and the water hydration in the asparagine gate region along the 1D minimum free energy path (black line) is clearly evident. The statistical errors of the 2D-PMFs in (A), (B) and (D) are in the range ~ 0.1-3 kcal/mol. The statistical errors of the 1D-PMFs in (C) are in the range ~ 0.1-1 kcal/mol. In all the plots, the positions of E286, PRD*a*_3_, PRA*a*_3_, D132, N139, S200 and S201 are labeled with text boxes.

The 1D PMFs traced out along the minimum free energy pathways (Fig. 2 C), which should describe the dominant activated reactive energetics for the water-mediated transport of the pumped protons in the P_M_’ and P_R_ states, also reveal several important findings. First, the free energy minimum at the PRA*a*_3_ in the pumping PMF of the P_R_ state suggests that PRA*a*_3_ is the major PLS, in agreement with the conclusions of ref (29, 30). It is interesting to note that in *ba*_3_-type C*c*O, PRA*a*_3_ is also suggested to be the PLS (31). However, PRD*a*_3_ is also actively involved in the pumping process since it shuttles the proton from E286 to the PLS (breaking its salt bridge to R481 and hydrogen bond with W172), as suggested by Wikstrom et al (32) based on non-reactive classical MD simulations. The participation of PRD*a*_3_ in the PT from E286 to PLS was also discussed by Yamashita et al. (17) in a reduced model for the C*c*O system. Second, PT from E286 to the PLS is thermodynamically unfavorable by ~ 8 kcal/mol in P_M_’ state, but favorable by 3 kcal/mol in P_R_ state (Fig. 2 C). Thus, ET to the BNC provides a thermodynamic driving force for PT to the PLS. Combined with the experimental results showing that ET is not complete without PT to the PLS in the E286Q mutant (33), one might conclude that ET to the BNC and PT to the PLS are coupled, each driving the other to its thermodynamically favored state.

### Transport of the Chemical Proton

Focusing next on the transfer of the chemical proton, we start from the low hydration state of the HC and calculate 1D PMFs for PT from E286 to the BNC in P_M_’ and P_R_ states, both with and without the PLS (PRA*a*_3_) protonated. The low hydration state is chosen for several reasons. First, the PT pathway is roughly horizontal to the membrane such that PRD*a*_3_ remains deprotonated throughout the PT process, and never rotates down to increase the hydration level in the HC. Second, classical MD simulations starting from the high hydration state and PRA*a*_3_ protonated relax to low hydration state within 30 ns once E286 is also protonated (i.e., following E286 reprotonation from the D channel, which is fast as further described below). The PMFs (*SI Appendix*, Fig. S3) show that PT from E286 to the Cu_B_ bound hydroxide in the BNC is thermodynamically unfavorable by more than 10 kcal/mol in the P_M_’ state, but favorable by ~ 5 kcal/mol in the P_R_ state. This suggests that ET from heme *a* to BNC also provides a thermodynamic driving force for the PT from E286 to the BNC (i.e., the formation of the F state) that leads to the chemical reaction. Interestingly, this conclusion is independent of whether the PLS is loaded (protonated) or unloaded (deprotonated) (*SI Appendix*, Fig. S3 E), and also is not sensitive to the hydration level in the HC (*SI Appendix*, Fig. S3 A-D).

### Rates of the Pumped and Chemical Proton Transport Events

Analysis of the rates (Table 1; reported as time constants, i.e., the inverse of rate constants) for PT to the PLS and BNC in the P_M_’ and P_R_ states provides deeper mechanistic insight into the proton pumping mechanism during the A→ P_R_→ F transition. In the P_M_’ state (before ET), PT from E286 to the PLS is thermodynamically unfavorable, but still faster than the experimental A**→**P_R_ transition rate (50 μs for *Rhodobactor sphaeroides* (8) and ~25 μs for *Paracoccus denitrificans* (34)). However, PT from E286 to the Cu_B_ bound hydroxide in the BNC is significantly slower. Thus, before ET it is kinetically prohibitive to transfer the chemical proton from E286 to the BNC, which would short-circuit pumping if it were to happen before PT to the PLS (25). Although forward PT from E286 to PLS is kinetically possible in the P_M_’ state, it is energetically unfavorable. Moreover, the reverse PT (PLS to E286) is even faster and outcompetes reprotonation of E286 through D-channel (Table 2). Thus, PT to the PLS is minimal before the ET. In contrast, after ET (in the P_R_ state) proton back flow from the PLS to deprotonated E286 is slower than the reprotonation of E286 through the D-channel. This allows timely reprotonation of E286 and prevents the proton loaded at PLS from leaking back to E286 and subsequently being consumed at the BNC. Thus, full loading of the PLS is achieved only after or concurrently with ET. In line with this, we note that the PT to PLS is still faster than PT to the BNC after the ET (Table 1, P_R_ state). This again prevents the above-mentioned short-circuiting and ensures that for the entire PT/ET process during A**→**F transition, regardless of the redox states of heme *a* and the BNC, PT to the PLS is not short-circuited by PT to the BNC.

**Table 1.** Calculated time constants (inverse of rate constants*) for PT of the pumped (E286→PLS), chemical (E286→BNC), and back leaked (PLS→E286) protons in the P_M_’ and P_R_ states, compared with experimental time constants for A→P_R_ and P_R_ →F transitions (8,34). The PT from E286 to BNC in P_R_ state with PLS protonated is the most physically relevant for the P_R_→F transition, and is in quantitative agreement with the experimental time constant. (See main text)

| State | E286→PLS ($\mu s$)<sup>#</sup> | E286→BNC ($\mu s$)<sup>#</sup> | | PLS→E286 ($\mu s$) |
| --- | --- | --- | --- | --- |
|  |  | Deprotonated PLS | Protonated PLS |  |
| $P_M'$ | $(2.6 \pm 0.2) \times 10^{-1}$ | $(4 \pm 3) \times 10^6$ | $(7.7 \pm 0.2) \times 10^8$ | $(1.5 \pm 0.4) \times 10^{-5}$ |
| $P_R$ | $2.1 \pm 0.3$ | $(3 \pm 2) \times 10^2$ | $(1.7 \pm 0.9) \times 10^2$ | $(1.2 \pm 0.6) \times 10^3$ |
| Exp ($A \rightarrow P_R$) | | 25-50 | | |
| Exp ($P_R \rightarrow F$) | | 200 | | |

* See SI for more information on the rate constant calculations. Errors are shown in parenthesis.

^#^For E286→PLS, the initial state has E286 protonated and the PLS deprotonated while the final state has E286 deprotonated and the PLS protonated, and vice versa for PLS→E286. For E286→BNC, the initial state has E286 protonated and a hydroxide bound to Cu_B_ in the BNC while the final state has E286 deprotonated and a water bound to Cu_B_ in the BNC.

**Table 2.**
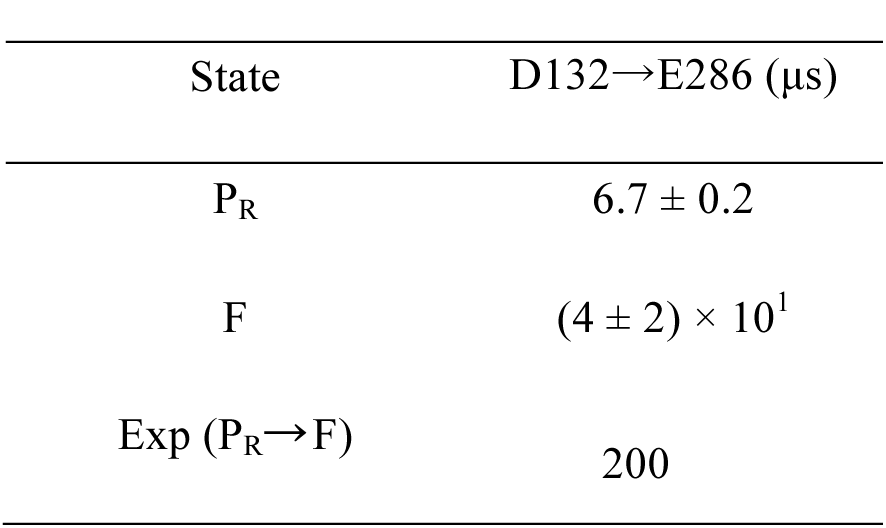
Calculated time constants (inverse of the rate constants*) for PT in the D-channel from protonated D132 to deprotonated E286 in the P_R_ and F states, compared with the experimental time constant for the P_R_ →F transition (8). The PLS is protonated in the simulation.

*See SI for more information on the rate constant calculations. Errors are shown in parenthesis.

Transfer of the chemical proton from E286 to the BNC is much faster after ET occurs because the electrostatic repulsion from the BNC is smaller in the P_R_ state than it is in the P_M_’ state (Table 1), reducing the PT free energy barrier (*SI Appendix*, Fig. S3 E). This supports the general expectation and our conclusion that ET to BNC facilitates the transfer of chemical proton to the catalytic site. Our calculated rate for transfer of the chemical proton in P_R_ state is in quantitative agreement with the experimental P_R_**→** F transition rate (8, 9), and is slower than the D-channel PT rates (see Table 2). Therefore, we conclude that the PT from E286 to the Cu_B_ bound hydroxide in the BNC is the rate-limiting step for the P_R_**→** F transition. To summarize, the above thermodynamic and kinetic results suggest that ET from heme *a* to BNC provides a thermodynamic driving force for PT both from E286 to the PLS *and* from E286 to the BNC, while PT from E286 to the BNC is the rate-limiting step for the P_R_**→** F transition.

### Comparison to Proposed Mechanisms

Our thermodynamic and kinetic results build upon a the previously proposed mechanism based on the orientation and connectivity of water chains in the HC (9, 19, 23, 25). In this mechanism, PT from E286 to the PLS and ET from heme *a* to the BNC are tightly coupled to each other and occur in concerted fashion during the A→P_R_ transition. The short-circuiting (i.e., premature PT to BNC before PT to PLS) is avoided by the absence of a water chain leading from protonated E286 to BNC in the P_M_’ state. However, this mechanistic proposal was based on classical nonreactive MD simulations in the absence of a shuttling excess proton. Our results support the conclusion that PT to the BNC is unfavourable in P_M_’ state, but given our treatment of the explicit proton transport they further explain why PT to the PLS is preferred after ET occurs, a question raised but not fully answered in ref (19).

Our results are also in partial agreement with the mechanism proposed by Faxen et al (8) in which PT from E286 to the PLS occurs after the formation of the P_R_ state, such that the ET and PT events are *sequential*. This mechanism was further discussed later by Lepp et al (35), who concluded that the electrometric signal during A→P_R_ transition is caused by the upward movement of K362 side chain rather than PT from E286 to PLS. However, it has been shown that this electrometric signal is large enough to include both the lysine swing and the PT from E286 to PLS. (19) Moreover, the electrometric signal was shown to be lost in the E286Q mutant (33), resulting in a product that was actually no longer pure P_R_, but a P_R_/P_M_’ mixture with the latter dominating (see Section 5.2. and Fig. 10 in ref. (14)). Based on these experimental findings, we suggest that the coupled PT/ET mechanism is at present the best explanation. (19) The results presented herein clearly show that ET drives PT from E286 to PLS, but they do not directly determine whether the PT/ET mechanisms are coupled or sequential. (8, 9)

### Proton Transport through the D-Channel

The D-channel is responsible for transporting two protons from the N-side of the membrane to E286 during the A→P_R_→F transition, one for pumping and the other for the chemical reaction. Three highly conserved asparagine residues, N139, N121, and N207, reside roughly one third of the way into the D-channel and form a constricted region (called the asparagine gate) (36). Previous nonreactive classical MD simulations have revealed a gating motion of N139 that controls the hydration state of D-channel (37, 38). However, that work hypothesized that PT through the D-channel follows a sequence in which the proton waits on the proton donor (D132) until a “water wire” is formed, then rapidly dissociates and transports through the pre-formed water wire to the proton acceptor (E286). As discussed above and in ref (28), this historically popular depiction of PT in aqueous systems is both misleading and inconsistent with the dynamically coupled and cooperative nature of the hydration environment and the excess proton migration.

Here and as we did earlier for the PT in the HC, we investigate how the motion of the excess proton is coupled to the change of the hydration level across the asparagine gate by explicitly calculating 2D PMFs in P_R_ and F states during the A**→** F transition (Fig. 2 D, *SI Appendix*, Fig. S4 A and B). In these PMFs the progress of the excess proton CEC through the D-channel (horizontal axis) and the degree of hydration of the asparagine gate region (vertical axis) are used as the two collective coordinates. The minimum free energy pathways are also identified on the 2D PMFs (black lines) and the corresponding 1D PMFs along the minimum free energy pathways (reaction coordinate) are plotted in *SI Appendix*, Fig. S4 C.

The D-channel PT process starts with protonated D132 and deprotonated E286. Initially, when D132 is protonated the space between N139 and N121 is narrow and dehydrated, forming an effective gate for PT (*SI Appendix*, Fig. S6 A). As the excess proton transitions to the water above D132 and approaches this gate, the N139 side chain rotates and opens a pathway for solvation and PT past the asparagine residues (*SI Appendix*, Fig S6 B). The transition state is reached when the excess proton is in the middle of the asparagine gate (*SI Appendix*, Fig S6 C). Once the excess proton shuttles through, the asparagine gate gradually closes and becomes dehydrated again (*SI Appendix*, Fig. S6 D). The curvy and non-horizontal nature of the minimum free energy pathways on the 2D PMF again indicates that PT and hydration changes are concerted and coupled processes. After traversing the gate region, the excess proton proceeds to the serine zone (S200 and S201), where it forms hydrogen bonds with the hydroxyl groups of the pore lining serine residues in a metastable state. Subsequently, the excess proton protonates E286 rotated in the “down” conformation (x≃5 Å *SI Appendix*, Fig. S4 C), reaching the global free energy minimum. Then, protonated E286 rotates up for either pumping or the chemical reaction (x≃1 Å *SI Appendix*, Fig. S4 C). The protonated E286 is slightly more energetically favorable in the P_R_ state than in the F states (*SI Appendix*, Fig. S4 C), likely due to the more negative charge on the BNC of the P_R_ state.

The 1D PMFs along the minimum free energy pathways on the 2D PMFs reveal free energy barriers for the proton to pass through the asparagine gate in the D-channel (*SI Appendix*, Fig. S4 C). For both states the calculated rates are much faster than the overall P_R_→F transition rate (Table 2), confirming that forward PT (D132 to E286) through the D-channel is not rate limiting in the P_R_→F transition. The D-channel PT rates are also faster than the PT back flow rates from the PLS, as discussed above, allowing for the fast reprotonation of E286 and preventing proton back flow from the PLS to deprotonated E286. It is plausible, based on these results, that the decoupling mutants, such as N139T (35) and N139D (39), disable kinetic gating by either slowing down PT through the D-channel (enabling back flow from the PLS to E286 to outcompete the reprotonation of E286; ref (14, 31, 38), or by slowing down the pumping rate (bypassing pumping altogether).

In recent years an alternative mechanism has been suggested (13, 20, 40), in which the excess proton approaches a protonated (neutral) E286 through the D-channel and facilitates proton pumping through a positively charged, biprotonated E286 transition state. In ref (20) the biprotonated mechanism was proposed because the proton pumping free energy barrier from singly protonated E286 calculated with the QM/MM computational methodology in that paper was too high. However, our results show that the approximate SCC-DFTB method used in ref (20) significantly overestimates the pK_a_ for aspartic acid deprotonation in bulk water and that the biprotonated E286 transition state is highly energetically unfavorable (see *SI Appendix*). These results suggest that the prohibitive pumping barrier reported in ref (20) could, at least in part, be an artifact of the SCC-DFTB method due to its overestimation of the proton affinity of E286 (see *SI Appendix* for further discussion).

## Conclusions

The quantitative multiscale reactive MD computational analysis of the explicit PT steps in C*c*O presented in this work, combined with previous experimental findings [see refs (14, 19, 21) and references therein], lead us to the following conclusions regarding the most likely sequence of PT and ET events during the A→P_R_→F transition. First, an electron is transferred from heme *a* to the BNC during the 25-50 μs A→P_R_ transition, likely following the chemistry that occurs at the BNC (33, 41). Either coupled with this ET event (during the A→P_R_ transition) or immediately after it (during the P_R_→F transition), a proton is transferred from a singly protonated E286 to the PLS (PRA*a*_3_). Experimental findings suggest that the former (coupled PT/ET) is more likely. (14, 19, 33) This PT induces and is accompanied by an increase in the HC hydration level from approximately four to eight waters. Second, E286 is rapidly reprotonated through the D-channel, and the HC relaxes back to the low hydration state. Third, the uncompensated negative charge in the BNC caused by ET triggers transfer of the chemical proton from E286 to the Cu_B_-bound hydroxide in the BNC, forming a water molecule. Fourth, E286 is reprotonated again through D-channel, accompanied by partial ET from Cu_A_ to heme *a* and proton ejection from PLS to the P-side.

Based on the above reaction sequence, we combine the PMFs and available experimental data to derive the free energy diagram for the A→F transition (Fig. 3). When the ET is allowed to the BNC without the possibility of protonating PLS (E286Q mutant), only 30% of P_R_ is formed, the rest remains as P_M_’ (ref (14), Fig. 10). From this, one can estimate that the ET from heme *a* to BNC alone (without protonation of PLS or BNC) is endergonic by ~ 0.5 kcal/mol. Therefore, starting from the P_M_’ state (state I, 0 kcal/mol), the PT from E286 to the PLS coupled with ET from heme *a* to heme *a3* leads to P_R_ state (state II) with energy level of −2.9 + 0.5 = −2.4 kcal/mol. Following this, assuming the pK_a_ of D132 is similar to that in bulk [which is ~ 3.9 (42)], the proton uptake of D132 from N-side bulk leads to state III (1.9 kcal/mol). Then the proton is transported from D132 to E286, forming state IV (−3.7 kcal/mol). The proton on E286 is subsequently transferred to the BNC, forming the F state (state V, −8.9 kcal/mol). The subsequent proton uptake of D132 from N-side bulk leads to state VI (−4.6 kcal/mol), followed by the second reprotonation of E286 that leads to state VII (−7.7 kcal/mol). Thus, proton release from the PLS to the P-side bulk is estimated to have exergonicity of 5 kcal/mol, completing the A→F transition and leading to state VIII (−12.7 kcal/mol). This 12.7 kcal/mol exergonicity reflects the overall driving force for the A→F transition, as estimated by experimental measurement. (14, 29)

**Fig. 3.**
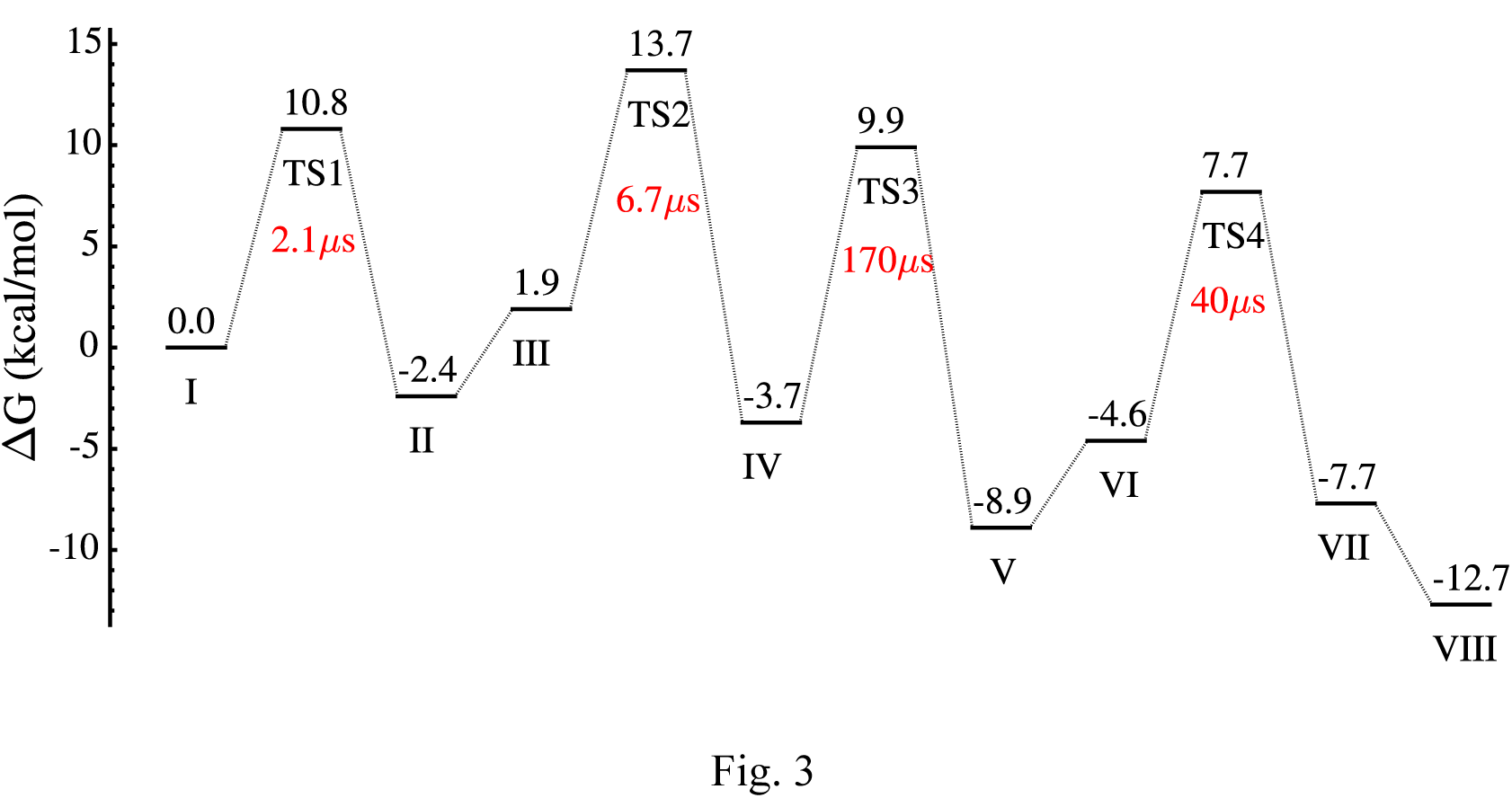
Free energy diagram for the reaction sequence during the A→F transition. The resting states are described in the main text. The transition states between them are labeled with “TS”, and the activation barriers are obtained from the free energy profiles (PMFs) in Figs. 2 C, S3 E, and S4 C. The black numbers indicates the energy levels for each state in kcal/mol. The red numbers indicate the time constants for the forward transitions between the two neighboring resting states.

Our results show that ET from heme *a* to the BNC drives both PT events (E286 to the PLS *and* E286 to the BNC). Among all of these processes, transfer of the chemical proton from E286 to the BNC is rate-limiting and our calculated rate is in quantitative agreement with experiment (8, 9). In this mechanism, the pathways that would decouple pumping from transfer of the chemical proton or damage the directionality of proton flow are avoided by kinetic gating in two ways. First, the fast PT from E286 to the PLS precedes proton transfer to the BNC during the entire A→F transition. Second, the possible proton back flow from the PLS to deprotonated E286 is avoided by fast reprotonation of E286 through the D-pathway combined with a large barrier for proton back flow from the PLS to E286 after ET from heme *a* to BNC (P_R_ state). Although PT in the D-channel is clearly coupled with solvation changes through the asparagine gate, it is never found to be rate limiting in the A→F transition. The multiscale reactive MD simulations presented herein have thus contributed to a comprehensive understanding of the functional mechanism of C*c*O, the important final enzyme in the respiratory electron transfer chain. Based on the success of this approach for CcO, we are optimistic that it can also be fruitfully applied to other redox-coupled proton pumping proteins.

## Material and Methods

The full structure of CcO from *Rhodobactor sphaeroides* (PDB code 1M56 (43)) was embedded in a dimyristoylphosphatidylcholine (DMPC) lipid bilayer and solvated by water molecules on each side of the membrane. MS-RMD simulations using metadynamics (MTD) (44) were performed to identify the PT pathways (45) in both the D-channel and HC. The FitRMD method (6, 7) was used to parameterize the MS-RMD models from QM/MM data for protonatable sites in CcO. The MS-RMD umbrella sampling calculation in the HC and D-channel were carried out by restraining 1) the excess proton CEC position (see *SI Appendix* for definition) along the PT pathway defined from the MTD procedure, and 2) the water density in a predefined box (see ref (28) for definition). Further details on simulation methods are provided in the *SI Appendix*.

## ACKNOWLEDGMENTS.

This research was supported by National Institutes of Health Grant R01-GM053148 (GAV, JMJS, RL, YP) and the Sigrid Jusélius Foundation (MW). The researchers used computing facilities provided by the Extreme Science and Engineering Discovery Environment (XSEDE), which is supported by National Science Foundation Grant OCI-1053575, as well as from the University of Chicago Research Computing Center (RCC), the Texas Advanced Computing Center at the University of Texas at Austin, and the U.S. Department of Defense (DOD) High Performance Computing Modernization Program. We also acknowledge Prof. Peter Brzezinski and Prof. Robert Gennis for helpful discussions, and Dr. Vivek Sharma, Prof. Ville Kaila, and Prof. Qiang Cui for providing equilibrated structures from their published work, which were helpful in determining the correct starting hydration state of the HC for our 2D PMFs. We also acknowledge Dr. J. G. Nelson for the initial simulation setup and helpful discussions in the early stages of this project.

## References

1. Knight C & Voth GA (2012) The Curious Case of the Hydrated Proton. Acc. Chem. Res. 45(1):101–109.

2. Knight C, Lindberg GE, & Voth GA (2012) Multiscale reactive molecular dynamics. J. Chem. Phys. 137(22).

3. Swanson JMJ, et al. (2007) Proton solvation and transport in aqueous and biomolecular systems: Insights from computer simulations. J. Phys. Chem. B 111(17):4300–4314.

4. Liang R, Li H, Swanson JMJ, & Voth GA (2014) Multiscale simulation reveals a multifaceted mechanism of proton permeation through the influenza A M2 proton channel. Proc. Natl. Acad. Sci. U. S. A. 111(26):9396–9401.

5. Yamashita T, Peng Y, Knight C, & Voth GA (2012) Computationally Efficient Multiconfigurational Reactive Molecular Dynamics. J. Chem. Theory Comput. 8(12):4863–4875.

6. Nelson JG, Peng Y, Silverstein DW, & Swanson JMJ (2014) Multiscale Reactive Molecular Dynamics for Absolute pK(a) Predictions and Amino Acid Deprotonation. J. Chem. Theory Comput. 10(7):2729–2737.

7. Lee S, Liang R, Voth GA, & Swanson JMJ (2016) Computationally Efficient Multiscale Reactive Molecular Dynamics to Describe Amino Acid Deprotonation in Proteins. J. Chem. Theory Comput. (In Press):DOI: 10.1021/acs.jctc.1025b01109.

8. Faxen K, Gilderson G, Adelroth P, & Brzezinski P (2005) A mechanistic principle for proton pumping by cytochrome c oxidase. Nature 437(7056):286–289.

9. Belevich I, Verkhovsky MI, & Wikstrom M (2006) Proton-coupled electron transfer drives the proton pump of cytochrome c oxidase. Nature 440(7085):829–832.

10. Kim YC, Wikstrom M, & Hummer G (2007) Kinetic models of redox-coupled proton pumping. Proc. Natl. Acad. Sci. U. S. A. 104(7):2169–2174.

11. Belevich I, Bloch DA, Belevich N, Wikstrom M, & Verkhovsky MI (2007) Exploring the proton pump mechanism of cytochrome c oxidase in real time. Proc. Natl. Acad. Sci. U. S. A. 104(8):2685–2690.

12. Kim YC, Wikstrom M, & Hummer G (2009) Kinetic gating of the proton pump in cytochrome c oxidase. Proc. Natl. Acad. Sci. U. S. A. 106(33):13707–13712.

13. Siegbahn PEM & Blomberg MRA (2010) Quantum Chemical Studies of Proton-Coupled Electron Transfer in Metalloenzymes. Chem. Rev. 110(12):7040–7061.

14. Kaila VR, Verkhovsky MI, & Wikstrom M (2010) Proton-Coupled Electron Transfer in Cytochrome Oxidase. Chem. Rev. 110:7062–7081.

15. Hammes-Schiffer S & Stuchebrukhov AA (2010) Theory of Coupled Electron and Proton Transfer Reactions. Chem. Rev. 110(12):6939–6960.

16. Kim YC & Hummer G (2012) Proton-pumping mechanism of cytochrome c oxidase: A kinetic master-equation approach. Biochim. Biophys. Acta 1817(4):526–536.

17. Yamashita T & Voth GA (2012) Insights into the mechanism of proton transport in cytochrome c oxidase. J. Am. Chem. Soc. 134(2):1147–1152.

18. Lu J & Gunner MR (2014) Characterizing the proton loading site in cytochrome c oxidase. Proc. Natl. Acad. Sci. U. S. A. 111(34):12414–12419.

19. Wikström M, Sharma V, Kaila VRI, Hosler JP, & Hummer G (2015) New Perspectives on Proton Pumping in Cellular Respiration. Chem. Rev. 115(5):2196–2221.

20. Goyal P, Yang S, & Cui Q (2015) Microscopic basis for kinetic gating in cytochrome c oxidase: insights from QM/MM analysis. Chem. Sci. 6(1):826–841.

21. Lepp H, Svahn E, Faxen K, & Brzezinski P (2008) Charge transfer in the K proton pathway linked to electron transfer to the catalytic site in cytochrome c oxidase. Biochemistry 47(17):4929–4935.

22. Riistama S, et al. (1997) Bound water in the proton translocation mechanism of the haem-copper oxidases. FEBS Lett. 414(2):275–280.

23. Wikstrom M, Verkhovsky MI, & Hummer G (2003) Water-gated mechanism of proton translocation by cytochrome c oxidase. Biochim. Biophys. Acta 1604(2):61–65.

24. Tuukkanen A, Kaila VRI, Laakkonen L, Hummer G, & Wikstrom M (2007) Dynamics of the glutamic acid 242 side chain in cytochrome c oxidase. Biochim. Biophys. Acta 1767(9):1102–1106.

25. Sharma V, Enkavi G, Vattulainen I, Róg T, & Wikström M (2015) Proton-coupled electron transfer and the role of water molecules in proton pumping by cytochrome c oxidase. Proc. Natl. Acad. Sci. U. S. A. 112(7):2040–2045.

26. Ghosh N, Prat-Resina X, Gunner MR, & Cui Q (2009) Microscopic pKa Analysis of Glu286 in Cytochrome c Oxidase (Rhodobacter sphaeroides): Toward a Calibrated Molecular Model. Biochemistry 48:2468–2485.

27. Goyal P, Lu J, Yang S, Gunner MR, & Cui Q (2013) Changing hydration level in an internal cavity modulates the proton affinity of a key glutamate in cytochrome c oxidase. Proc. Natl. Acad. Sci. U. S. A. 110(47):18886–18891.

28. Peng Y, Swanson JMJ, Kang S-g, Zhou R, & Voth GA (2015) Hydrated Excess Protons Can Create Their Own Water Wires. J. Phys. Chem. B 119(29):9212–9218.

29. Wikstrom M & Verkhovsky MI (2007) Mechanism and energetics of proton translocation by the respiratory heme-copper oxidases. Biochim. Biophys. Acta 1767(10):1200–1214.

30. Lee HJ, Ojemyr L, Vakkasoglu A, Brzezinski P, & Gennis RB (2009) Properties of Arg481 Mutants of the aa(3)-Type Cytochrome c Oxidase from Rhodobacter sphaeroides Suggest That neither R481 nor the Nearby D-Propionate of Heme a(3) Is Likely To Be the Proton Loading Site of the Proton Pump. Biochemistry 48(30):7123–7131.

31. Chang HY, et al. (2012) Exploring the proton pump and exit pathway for pumped protons in cytochrome ba(3) from Thermus thermophilus. Proc. Natl. Acad. Sci. U. S. A. 109(14):5259–5264.

32. Wikstrom M, et al. (2005) Gating of proton and water transfer in the respiratory enzyme cytochrome c oxidase. Proc. Natl. Acad. Sci. U. S. A. 102(30):10478–10481.

33. Gorbikova EA, Belevich I, Wikstrom M, & Verkhovsky MI (2008) The proton donor for O-O bond scission by cytochrome c oxidase. Proc. Natl. Acad. Sci. U. S. A. 105(31):10733–10737.

34. Belevich I, et al. (2010) Initiation of the proton pump of cytochrome c oxidase. Proc. Natl. Acad. Sci. U. S. A. 107(43):18469–18474.

35. Lepp H, Salomonsson L, Zhu JP, Gennis RB, & Brzezinski P (2008) Impaired proton pumping in cytochrome c oxidase upon structural alteration of the D pathway. Biochim. Biophys. Acta 1777(7-8):897–903.

36. Han D, et al. (2006) Replacing Asn207 by Aspartate at the Neck of the D Channel in the aa3-Type Cytochrome c Oxidase from Rhodobacter sphaeroides Results in Decoupling the Proton Pump. Biochemistry 45(47):14064–14074.

37. Henry RM, Yu CH, Rodinger T, & Pomes R (2009) Functional Hydration an Conformational Gating of Proton Uptake in Cytochrome c Oxidase. J. Mol. Bio 387(5):1165–1185.

38. Henry RM, Caplan D, Fadda E, & Pomes R (2011) Molecular basis of proto uptake in single and double mutants of cytochrome c oxidase. J Phys Conden Matter 23(23):234102.

39. Namslauer A, Pawatet AS, Gennis R, & Brzezinski P (2003) Redox-couple proton translocation in biological systems: Proton shuttling in cytochrome oxidase. Proc. Natl. Acad. Sci. U. S. A. 100(26):15543–15547.

40. Siegbahn PEM & Blomberg MRA (2008) Proton Pumping Mechanism i Cytochrome c Oxidase. J. Phys. Chem. A 112(50):12772–12780.

41. Karpefors M, Adelroth P, Namslauer A, Zhen YJ, & Brzezinski P (2000) Formation of the "peroxy" intermediate in cytochrome c oxidase is associated wit internal proton/hydrogen transfer. Biochemistry 39(47):14664–14669.

42. Lide DR (2004) CRC handbook of chemistry and physics.

43. Svensson-Ek M, et al. (2002) The X-ray Crystal Structures of Wild-type and EQ(286) Mutant Cytochrome c Oxidases from Rhodobacter sphaeroides. J. Mol. Bio 321(2):329–339.

44. Laio A & Parrinello M (2002) Escaping free-energy minima. Proc. Natl. Acad. Sci. U. S. A. 99(20):12562–12566.

45. Zhang Y & Voth GA (2011) Combined Metadynamics and Umbrella Samplin Method for the Calculation of Ion Permeation Free Energy Profiles. J. Chem Theory Comput. 7(7):2277–2283.

